# Analog nitrogen sensing in *Escherichia coli* enables high fidelity information processing

**DOI:** 10.1101/015792

**Authors:** M. Komorowski, J. Schumacher, V. Behrends, T. Jetka, Mark H. Bennett, A. Ale, S. Filippi, J.W. Pinney, J.G. Bundy, M. Buck, M.P.H. Stumpf

## Abstract

The molecular reaction networks that coordinate the response of an organism to changing environmental conditions are central for survival and reproduction. *Escherchia coli* employs an accurate and flexible signalling system that is capable of processing ambient nitrogen availability rapidly and with high accuracy. Carefully orchestrated post-translational modifications of PII and the glutamine synthetase allow *E. coli* to trace nitrogen availability in a continuous, decidedly non-digital fashion. We measure the dynamic proteomic and metabolomic responses to trace the analog computations, and use an information theoretical framework to characterize the information capacity of *E. coli’*s nitrogen sensing network: we find that this system can transmit up to 9bits of information about the nitrogen state. This allows cells to respond rapidly and accurately even to small differences in metabolite concentrations.

## INTRODUCTION

Post-translational modifications (PTMs) play a pivotal role in cell signalling, allowing rapid responses through reversible modifications of proteins. Reversible covalent modifications at specific protein residues allow for rapid (and low-metabolic burden) signalling and regulation (Khoury *et al*, 2011). One of the best-studied bacterial PTM systems is the bicyclic nucleotidylation cascade of PII and glutamine synthetase (GS) that regulates responses to environmental nitrogen concentrations in *Escherichia coli*. Nitrogen-status-dependent uridylylation of PII controls GS adenylylation and activity (Fig. 1A) (van Heeswijk *et al*, 2013). Glutamine (GLN) signals nitrogen sufficiency through binding to the bifunctional uridylyl–transferase/removase (UT/UR), stimulating the de-uridylylation of PII-UMP. PII acts on the bifunctional adenylyl–transferase/removase (AT/AR) of GS, where PII stimulates the transfer and PII-UMP the removase functions of AT/AR. αKetoglutarate (αKG) binds to PII, changing its regulatory capacity so that the GS adenylylation state depends on both GLN and αKG, which together define the nitrogen status. In response to changes in the nitrogen status PII also regulates the two-component nitrogen regulators NtrB/NtrC (Bueno *et al*, 1985; Wedel *et al*, 1990), thereby directly controlling transcription of 45 genes in response to nitrogen starvation, including the gene coding for GS, *glnA* (Zimmer *et al*, 2000). While the whole system can be qualitatively understood on the basis of the regulatory connections (Fig. 1A), mathematical approaches are needed to allow us to quantitatively understand and model how the system responds to different nitrogen levels (Bruggeman *et al*, 2005; Yuan *et al*, 2009; Ma *et al*, 2009; Okano *et al*, 2010; Hart *et al*, 2011; Lodeiro & Melgarejo, 2008). Such models provide a mechanistic and testable description of the dynamics at the level of populations of cells.

**Figure 1:**
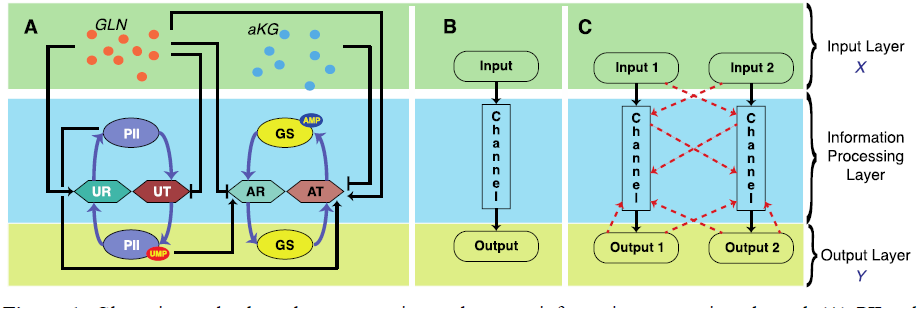
Glutamine and α-ketoglutarate sensing pathway as information processing channel. (**A**) PII and GS states are regulated by the bifunctional enzymes uridylyltransferase/uridylyl-removing enzyme (UT/UR) and adenylyltransferase/ adenylyl-removing enzyme (AT/AR), respectively. Inputs of the pathway, GLN and αKG, allosterically regulate AT/AR and UT/UR. The activity of AT/AR is also regulated by the signalling protein PII. Outputs of the pathway as uridylated PII and de-adenlylated GS indicate nitrogen scarcity. (**B**) Conventional view of the information channel, which connects one input with one output. (**C**) The pathway exhibits a complex connection pattern, where elements of the information processing layer interact with each other and with the outputs.

Studies that quantify information capacity of bacterial signalling circuits beyond chemotaxis (Kollmann *et al*, 2005; Endres & Wingreen, 2009) and quorum sensing (Mehta *et al*, 2009) are not available. Particularly, none of the PTM systems have been analysed in this context.

Experiments on mammalian single cells (Cheong *et al*, 2011; Uda *et al*, 2013) reveal that their signalling machinery is capable of transmitting the information needed to take binary decisions.

Quantitative measurements of modification states of signal transduction proteins are only rarely available. In particular, we lack experimental techniques to obtain measurements of metabolite changes and/or molecular noise at the level of single bacterial cells. Therefore, if we wish to investigate how molecular responses to environmental changes are orchestrated we either have to rely on partial single cell measurements of a few selected molecular protein species but without access to PTMs or metabolite abundances; or we use comprehensive population-level experimental measurements and analyse the resulting mechanistic model of the stress response. Here we opt for the latter and integrate population measurements with available knowledge regarding noise in similar systems to show that the bacterial PTM system has the potential to precisely and rapidly process information and to distinguish quantitatively different signals in order to infer the ambient nutrient availability

We use a combination of experimental methods to provide quantitative data to parameterize the models, including targeted proteomics and NMR analysis of metabolites. We directly measure the absolute amounts of PII, PII-UMP, GS, GS-AMP, GLN, and αKG in responses to perturbations in nitrogen availability *in vivo*, and use the data to probe the dynamics within this biological control system.

This molecular machinery allows the cell to process information in a way that can be cast in the language of Bayesian inference. We demonstrate that a direct relation between signalling and Bayesian statistical methodology provides a general conceptual, quantitative and computational framework for the analysis of information processing in biochemical circuits, and derive widely applicable insights into how biological organisms can achieve (near-)optimal signal transduction using solely molecular interactions and reactions.

## RESULTS

### Experimental Setup

The nitrogen status of *Escherichia coli* is predominantly given by the abundances of assimilated nitrogen in the form of glutamine (GLN), and of the Krebs cycle metabolite α-ketoglutarate (αKG). In order to capture the PTM changes of PII and GS in response to changes in nitrogen status, we grew wild-type *E. coli* (NCM3722) in batch culture in defined minimal media with an initial 3 mM NH_4_Cl concentration (Fig. 2A) (Schumacher *et al*, 2013). This concentration gives a nitrogen replete-condition at the beginning of growth, but as cells multiply the NH_4_^+^ is increasingly consumed, resulting in nitrogen limitation and then starvation (run out). Following 40 minutes of starvation, fresh NH_4_Cl (3 mM) is added (spike). We sampled at nine time points across this process (with the timing of the spike defining the t=0 time point) in order to capture the complete transition from NH_4_Cl-replete to NH_4_Cl starvation, and back to nitrogen-replete status (Fig. 2A), and then measured these samples for key metabolite and protein concentrations.

**Figure 2:**
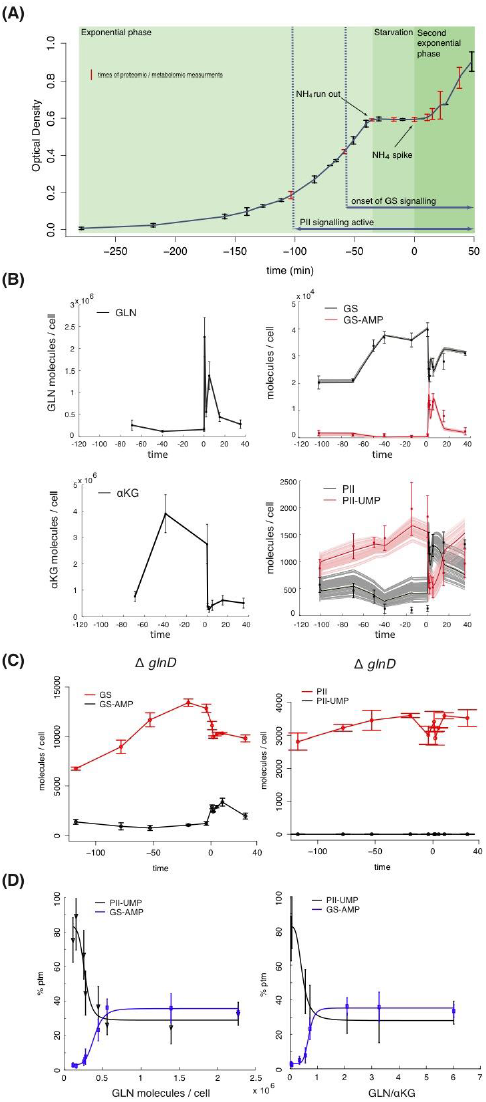
(**A**) The growth curves over 5 experiments show very little variation is the behaviour of the population dynamics. Error bars correspond to the standard error of the mean, and the red bars indicate time-points at which samples were taken for quantitative metabolomics and proteomics analysis (see *Supplementary Information* for more details). (**B**) Total abundances of the inputs (GLN, αKG) and outputs (PII, PII-UMP, GS, GS-AMP). (**C**) Total abundances of outputs in the UT/UR (*glnD*) knock-out. (**D**) PTM level of PII and GS as a function of the number if GLN molecules per cell (left) and of GLN/αKG ratio. Errors bars correspond to standard errors over three biological replicates, lines are the results from the simulations using parameters drawn from the inferred posterior distributions of the mathematical model of the pathway in Fig. 1A.

### Simultaneous measurements of metabolites and post-translational states of signalling proteins reveals sensitivity of the pathway

LC-MRM-MS can be used to directly measure targeted post-translational modifications, including phosphorylation, acetylation, methylation and ubiquitination (Picotti & Aebersold, 2012). The precise quantification of post-translational modifications in complex biological samples remains challenging for a combination of reasons, including the instability of modified peptides, incomplete protein extraction, incomplete proteolysis and artifactual protein modifications (Aebersold *et al*, 2013). Here we use a recently reported robust sample preparation protocol for LC-MRM-MS using labelled whole proteins as internal standards rather than individual peptides or polypeptides (protein standard absolute quantification, PSAQ), which to a large degree avoids these problems (Schumacher *et al*, 2013). We can directly and reproducibly measure the abundance of both the modified and un-modified PII and GS peptides (see *Supplementary Information*) and thus calculate the number of PII and GS molecules per cell (Fig. 2B). We also calculate the numbers of GLN and αKG molecules per cell based on NMR measurements. As expected (Ninfa *et al*, 2000; Yuan *et al*, 2009), PII uridylylation levels are inversely correlated with both intracellular GLN concentrations and the levels of GS adenylylation. To probe the model depicted in Figs. 1. and 3, and to capture the coupling of PII uridylylation with GS adenylylation *in vivo*, we generated a NCM3722ΔglnD knock-out mutant (*glnD* codes for UT/UR), which shows that when PII uridylylation is lost, GS adenylylation is no longer dependent on GLN levels (Fig. 2C). The rapid changes in protein PTMs, clearly related to metabolite changes, not only indicate the wide range of PII and GS PTM states that can be measured under our experimental conditions, but also demonstrate that our sampling and analysis pipeline provides high-quality data without substantial losses of PTMs.

**Figure 3:**
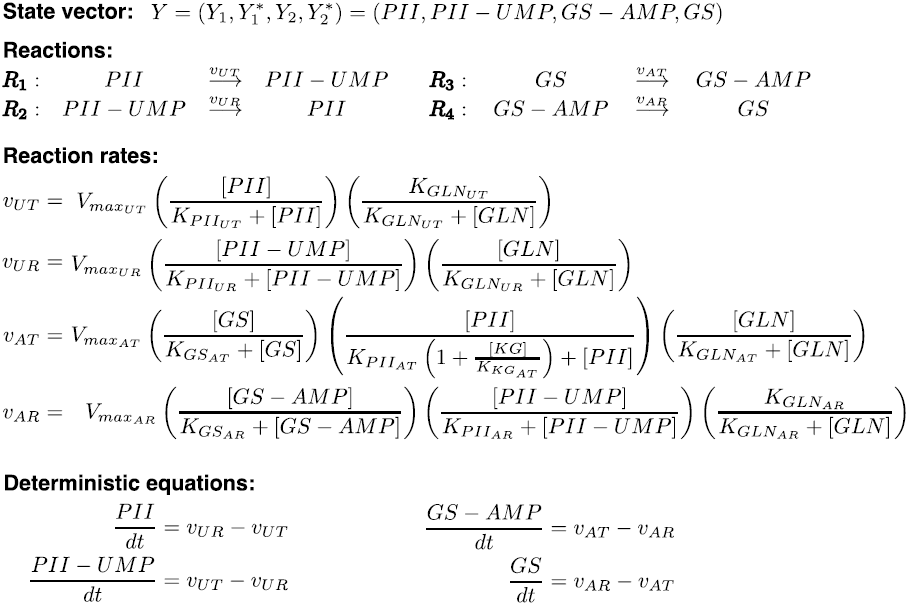
Mathematical description of reactions involved in the model depicted in the Fig. 1A together with ordinary differential equations describing model dynamics (see also reference (Yuan *et al*, 2009), main text and *Supplementary Information*.)

To determine the responsiveness of PII and GS PTMs to changes in GLN and αKG levels, we calculate the levels of PII and GS PTMs, taking into account the total PII and GS concentrations. As expected, for the constitutively transcribed *glnB* (*glnB* codes for PII), PII levels remain constant over the time course. GS levels increase by approximately 2.3 fold over the course of the experiments, as NH_4_ starvation induces *glnA* expression. The relative PII-UMP and GS-AMP levels are a function of the intracellular GLN concentration and the GLN/αKG ratio, since the fraction of PII-UMP should exclusively depend on GLN, whereas the relative GS-AMP levels depend on the GLN/αKG ratio. We observe an inverted sigmoidal relationship between GLN and PII-UMP levels (Fig. 2D), with a responsive range between 0.2 mM and 0.85 mM GLN (1.2×10^5^ – 5.1×10^5^ molecules per cell). GS-AMP is responsive at higher GLN concentrations (0.4 mM), with a clear sigmoidal dependency both with regards to GLN levels and the GLN/αKG ratio, providing evidence of an underlying cooperative mechanism of GS. In the NCM3722ΔglnD mutant, GLN sensing is uncoupled and these dependencies are lost; nevertheless we still find higher GS-AMP levels in samples with low αKG levels, indicating that in NCM3722ΔglnD, un-uridylylated PII bound to αKG can modulate the AT/AR activity (Fig. 2C).

The limited dynamic range of the PTMs as a function of GLN (or GLN/αKG ratio) is noteworthy: the relative PII-UMP and GS-AMP levels are confined to the 100%-30% and 0-35% ranges, respectively (Fig. 2D). Naively we might expect that precise sensing would exploit the whole range (Tkacik *et al*, 2009). Here, however, the same enzyme, UT/UR, carries out both uridylylation and deuridylylation of PII; equally, AT/AR adenylylates and deadenylylates GS. The steady states at high GLN concentrations are thus obtained dynamically by balancing the relative functionalities of the two enzymes. This dynamic tension allows rapid response to decreasing GLN availability (Kim *et al*, 2012): if GLN decreases deuridylylation and adenylation cease, and the fractions of uridylylated PII and de-adenylated GS increase rapidly because uridylylation and deadenylylation are already ongoing.

### The analog nature of nitrogen availability sensing

Direct comparison of the growth curve (Fig. 2A) with the modification states of the signalling proteins (Fig. 2B) indicates that the bicyclic system senses scarcity well before NH_4_^+^ depletion. Growth stops abruptly after the run-out of NH_4_^+^ and the onset of nitrogen starvation (-40 minutes in Fig. 2C), and we observe a relatively constant ratio of GLN/αKG; during this phase PII is mostly uridylylated and GS essentially de-adenylylated. At time 0, NH_4_Cl is introduced again into the medium, which results in the rapid depletion of αKG as nitrogen is assimilated, and a reversal in the abundances of PII–UMP and GS–AMP (Fig. 2B). Interestingly, within 30 seconds after the NH_4_^+^ spike, GLN levels consistently peak at their maximum (αKG levels drop by similar absolute amounts), probably due to the very high levels of fully active GS that is assimilating GLN (Okano *et al*, 2010) before GS becomes deactivated through adenylylation within roughly 1 minute following NH_4_^+^ addition. Several minutes post spike a dynamic steady state equilibrium is regained, and all molecules of the bicyclic system return towards the initial state (Hart *et al*, 2011).

*E. coli* growth experiments performed under nitrogen starvation show that the doubling time of the cells increases long before all of the available nitrogen as ammonium has been consumed (Fig. 2A). This growth characteristic together with the PTM dynamics clearly indicates that the amount of nitrogen available to the cell is not sensed in a simple binary or digital (PRESENT/ABSENT) manner but rather as an essentially continuous or analog signal. What has been unknown to date is how much information is or can be processed by such a system, i.e. how accurately can cells sense their nitrogen status *in vivo*.

Analog information processing can be more nuanced than digital information processing (Daniel *et al*, 2013). In particular, it tends to be easier to implement differentiation and integration of signals using analog logic building blocks; as a rule it is also less energy consuming (Sarpeshkar, 2014). Both of the above observations are also true for physical and engineering systems, even though digital computation has come to predominate primarily due to the availability of small, cheap and reliable electronic components. Thus analog information processing can be distinctly advantageous for detecting subtle variations in an environmental signal(Sarpeshkar, 1998; 2014).

### Probabilistic model of nitrogen sensing

When considering signal transduction processes we can make use of Shannon’s concept of a communication channel which links input and output (Fig. 1B) (Cover & Thomas, 2012). This simple relationship is difficult to reconcile with the more complex process depicted in Fig. 1A, where we have two inputs, GLN and αKG, and two outputs, uridylylated PII and non-adenylylated, active, GS. These outputs feed back to the protein complexes and molecules making up the information channels (*e.g*. uridylylated PII activates the adenylyl removase that deactivates adenylylated GS). Information flow in the nitrogen status sensing mechanism of *E. coli* thus more resembles the diagram shown in Fig. 1C, where inputs and outputs can affect the information processing (i.e. channels) directly and indirectly and in a variety of ways (Jiang *et al*, 1998; 2007).

In order to mathematically model the complexity of the information transmission process we consider the stimuli that signal availability of nitrogen, *X* = (*X*_1_,*X*_2_)*^T^*, where *X*_1_ and *X*_2_ denote GLN and αKG respectively, and which are jointly distributed according to probability distribution P(*X*_1_,*X*_2_). The levels of uridylylated PII-U and unmodified GS (as numbers of molecules), which signal nitrogen starvation, are denoted by *Y*_1_^*^ and *Y*_2_^*^, respectively. The forms that signal nitrogen abundance, unmodified PII and GS-A, are denoted by *Y*_1_ and *Y*_2_. The output of the system is therefore given as *Y* = (*Y*_1_,*Y*_1_^*^,*Y*_2_,*Y*_2_^*^)*^T^*. In order to capture the essentially probabilistic aspect of information transmission we considered the joint probability distribution over the abundances of *Y*_1_^*^ and *Y*_2_^*^, and thus represent the network performance in terms of the conditional probability, *P*(*Y*|*X*).

When considering fidelity of signal transduction processes the state of the art method (Cheong *et al*, 2011; Tkacik *et al*, 2008; Selimkhanov *et al*, 2014) is to reconstruct *P*(*Y*|*X*) based on single cell measurements. Alternatively a model of *P*(*Y*|*X*) can be used and the two approaches are complementary.

Single cell experiments usually induce additional variability resulting from fluorescence quantification, reporter copy number variation, perturbations introduced by reporter genes or variable antibody staining efficiency. Moreover, such experiments usually measure single input – single output relations and neglect many of the factors that contribute actively to cell to cell variability, which, for individual cells, may be means of adaptation and cannot be interpreted as noise. These may include differences in cell cycle progression, differences in auto- and para-crine signalling due to non-uniform distribution of cells on a culture-plate, variability in gap junction formation with the neighbouring cells etc. As a result such experiments are likely to underestimate fidelity of signal transduction systems.

These potential problems can largely be avoided by modelling the noise appropriately, It comes at the cost of analysing an idealized model of signalling pathway, where we have to make assumptions about the ways in which noise enters the dynamics. Both the direct experimental route and the model-based approaches have their own drawbacks and, given current experimental capabilities at the single cell level, both perspectives need to be pursued understand information flow processes in signal transduction pathways.

In order to examine the roles of PTMs in information processing we only have access to population measurements and therefore must use a model based approach as noise measurements are currently not obtainable at single cell level. In order to construct a model of *P*(*Y*|*X*) we first consider the functional relationship between nitrogen-related stimuli and the enzyme activities provided by solutions to equations that describe the temporal evolution of *Y* = (*Y*_1_,*Y*_1_^*^,*Y*_2_,*Y*_2_^*^)*^T^* (Fig. 3). Using the quantitative protein and metabolite data, we obtain estimates for the kinetic parameters that capture the functional relationships between GLN and αKG and their corresponding enzymes GS and PII. We use our Bayesian framework (Liepe *et al*, 2014; Toni *et al*, 2009) to estimate the parameters directly from the time-resolved metabolomic and proteomic data. Simulations of the deterministic model, shown by sampling parameters from the inferred posterior distributions (shown in Fig. Supp. 5), are in excellent agreement with the available data (Fig. 2B).

Given these parameter estimates we can calculate the probabilistic distribution of outputs given inputs, *P*(*Y*|*X*), as modelled by the chemical master equation. However, given the high number of PII and GS molecules involved (approximately 2300, and 20000-40000, respectively), we can also use the linear noise approximation (Van Kampen, 2011; Komorowski *et al*, 2009) to describe *P*(*Y*|*X*) without any significant loss of accuracy (Wallace *et al*, 2012). The probability distribution, *P*(*Y*|*X*), constructed in this way provides the stochastic version of the model in Fig. 3 (see also *Supplementary Information*). It allows us to describe the action of the information channel (Fig. 1B,C). In order to quantify its information capacity we develop a theoretical framework outlined below. We can thus quantify the mutual information (Cover & Thomas, 2012) – the canonical measure for the information shared between two variables – characteristic for *E. coli* nitrogen sensing.

### An Optimality Criterion for Information Transmission

In an information-theoretical context – including the situation considered above – the central quantity of interest is the amount of information that can flow through an information channel. Typically it is measured via Shannon’s mutual information (Cover & Thomas, 2012) between input, X, and output, Y,

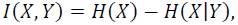
where *H*(*X*) and *H*(*X|Y*) are the entropy of the distribution over input *X*, and the average conditional entropy of *X* given Y, respectively. *I*(*X,Y*) is generally interpreted as the reduction in uncertainty about *X* once the value of *Y* is known; this is often also seen as a more general measure of correlation between two random variables *X* and *Y*. Importantly, *I*(*X, Y*) depends on matching between the *P*(*Y*|*X*) and *P*(*X*). The maximum amount of information that can flow through an information channel, *P*(*Y*|*X*), is known as *channel capacity* (Cover & Thomas, 2012)

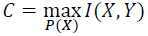

This measures the efficiency of a communication channel; in this case, the ability of *E. coli* to sense and respond to changes in ambient nitrogen availability. In order to calculate the maximal achievable *I*(*X, Y*), *i.e*. the channel capacity, we need to determine which input *P*(*X*) the system *P*(*Y*|*X*) is best adapted to.

We consider the conditional probability P(X|Y) as the inference that the cell draws about the environment (here as the ambient nitrogen abundances encoded by GLN and αKG). If this estimate of the input X is unbiased then its variance, 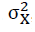, must obey 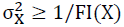, where *F1*(*X*) is the Fisher information (Brunel & Nadal, 1998; Komorowski *et al*, 2011) for given X,

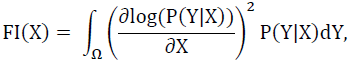
where Ω is the space of possible signals. In the asymptotic setting the variance of the estimate is given precisely by the inverse of the Fisher information, 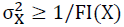.

Under very general conditions

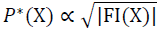
is the distribution that maximizes the mutual information, *I*(*X,Y*), between inputs, *X*, and outputs, *Y* and allows us to calculate the channel capacity Brunel & Nadal, 1998). In Bayesian inference this distribution is also known as the reference prior. This relationship nontrivially connects the mutual information and the Fisher information. Given the asymptotic interpretation 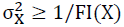, this criterion states that frequent inputs should be recognized/processed with high confidence (see below), whereas more rarely occurring signals (states of the random variable X) need not be inferred with similarly high accuracy. By balancing the frequency of inputs with the constraints of the system we find that the input distribution, *P*(*X*), that maximises the mutual information is given by FI(X), the Fisher Information associated with the inputs, X. In summary, the optimal distribution of inputs, *P*^*^(*X*), is defined in terms of uncertainty of inferences, P(X|Y), that the cell draws about the environment state, X. In the optimal scenario signals occur at a frequency that is proportional to the inverse of the uncertainty (which is measured as the standard deviation of the distribution P(X|Y)). In other words, common events and signals are sensed and processed with high precision, see Fig. 4. Signal transduction systems may thus evolve to ensure that the information transmission is maximized given the ambient inputs (Frank, 2009). This link between the mutual information and the Fisher information constitutes a general computational approach to analyse optimal information processing in biochemical signalling systems (Libby *et al*, 2007).

**Figure 4:**
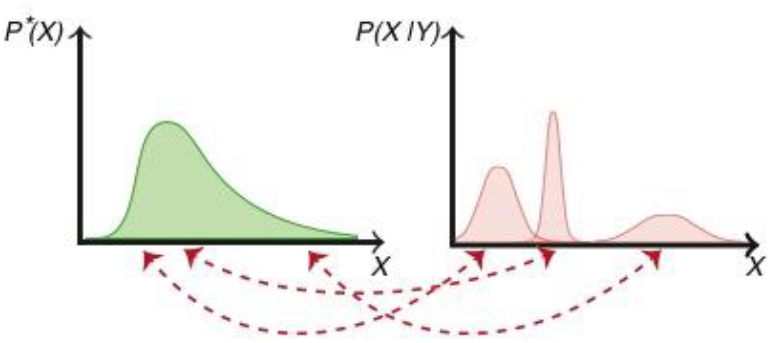
The optimal input distribution for a channel *P*^*^(*X*) is defined in terms of the uncertainties associated with the inferences of P(X|Y). For each intensity of the signal X, the optimal probability *P*^*^(*X*) is inversely proportional to the associated standard deviation of the distribution P(X|Y).

### Quantifying the information processing capacity of the nitrogen-sensing system of *E. coli*

In order to illustrate our information-theoretic perspective on signalling of nitrogen status we first consider the case of GLN sensed through PII (Yuan *et al*, 2009; van Heeswijk *et al*, 2013), the left-hand part of the system depicted in Fig. 1A; this conforms to the classical simple communication channel in Fig. 1B. In Fig. 5A we show the derived input-output relationships for GLN and PII-U, and we can derive the Fisher Information from the stochastic model (Komorowski *et al*, 2011; 2012), which provides us with the optimal input distribution (Fig. 5B). By sampling from the parameter posteriors obtained from the data, we then obtain a posterior over the channel capacity, which is centred around a capacity of C≈4.15 bits (Fig. 5C). The capacity, C, depends on kinetic rates that define P(Y|X). Averaging over posterior distributions takes into account uncertainty regarding these rates.

**Figure 5:**
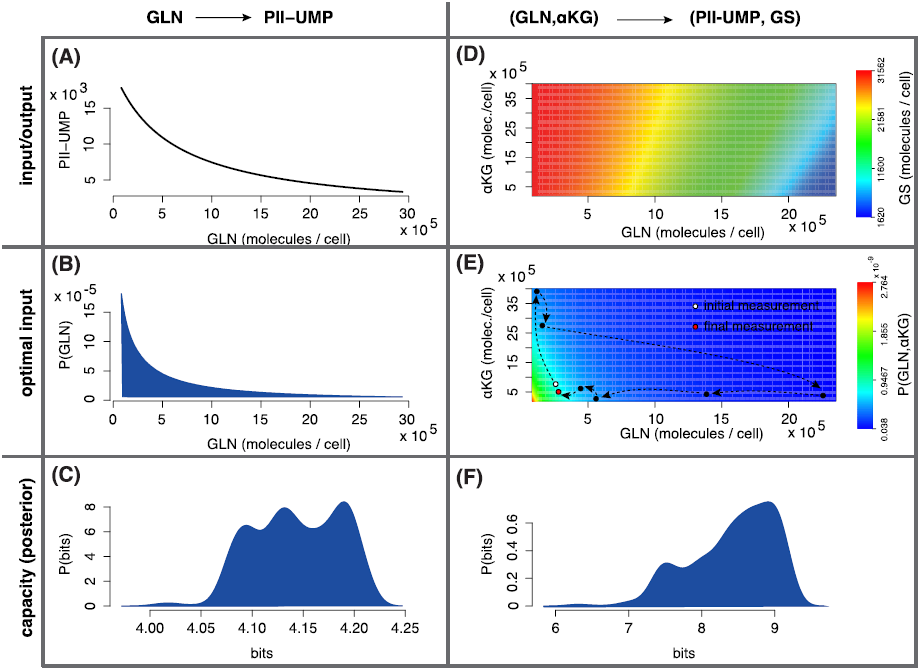
Optimal information processing via GLN – PII-U and (GLN, αKG) – (PII-U, GS) channels. (**A,D**) Input – output relations, calculated using the model in Fig. 2C, presented as linear plot (A) for GLN – PII-U and as heatmap (B) for (GLN, αKG) – (PII-U, GS). In the latter case unmodified GS is plotted. (**B,E**) Optimal input distributions calculated using the optimality criterium (Fig. 4). In addition to the heatmap, pannel (E) contains the same experimental measurements of GLN and αKG as in the Fig. 2B plotted as arrow connected black dots. (**C,F**) Posterior distributions of the information capacities C obtained from model parameter posterior (see also Fig. Supp. 6).

The above description captures only the information transmitted via the PII part of the nitrogen signalling system. In Fig. 5D-F we repeat the same analysis but consider both inputs simultaneously with their two corresponding outputs (Yuan *et al*, 2009; van Heeswijk *et al*, 2013), which is an example of the more complicated information processing network shown in Fig. 1C. The input-output relationship and optimal distribution are shown in Fig. 5D and Fig. 5E, respectively. By sampling from the parameter posterior, we find that the maximum amount of information that can be transmitted is approximately 9 bits. In light of recent estimates of the channel capacity in mammalian signal transduction (Cheong *et al*, 2011; Uda *et al*, 2013) this might seem surprisingly high. Our approach, however, incorporates from the outset a probabilistic model to describe the biochemical reactions, which allows us to directly describe the information content between inputs and outputs. Here, of course, we consider the system isolated from sources of noise that typically affect single cell data. Moreover, the high capacity is a result of the high abundances of GS and PII, as well as the dynamics of the joint signalling system, which is characterized by fast PTM processes, that are capable of tracing the nitrogen state with high accuracy.

Together with the optimality criterion discussed before, Figs. 5B and 5E reveal that the PTM system underlying nitrogen sensing in *E. coli* senses GLN and αKG with high fidelity at low abundances, and with reduced fidelity at high abundances. We would expect high levels of selection pressure for systems for sensing the availability and uptake of a key nutrient such as nitrogen. Intuitively, it makes sense that the information signalling system has been selected to give the highest precision for low levels of nutrient abundance, and that lower precision would suffice when ample levels of nutrients are available. In light of this we can revisit the results shown in Fig. 2: under low GLN conditions, sensing and response are high-precision as PII uridylylation and GS adenylylation change sensitively in this range. These results also suggest that one major physiological role of GS adenylylation is to finely tune nitrogen assimilation fluxes and not necessarily just to prevent a glutamate depletion upon a rapid ammonium upshift as previously proposed (Kustu *at al*, 1984).

Our study suggests that the nitrogen sensing PTM system (consisting of GS and PII and their associated (de)activating enzymes), as well as other PTM systems, may serve to perform reliable analogue information processing *in vivo*. The model, together with the optimality criterion derived here, explains how careful tuning and matching of signal levels to the sensing machinery allows molecular mechanisms to achieve high fidelity of signalling and information processing. The experimental and theoretical methodology presented here is general and can be readily transferred to study other signalling systems. In contrast to other studies of biological information processing, we used population data to reconstruct dynamics and noise characteristics of signalling, because technology to measure input-output relationships in a metabolite sensing systems at the level of single bacterial cells does not currently exist.

Our model assumes (i) that within our experimental conditions the PII homologue GlnK does not influence the adenylylation state of GS; and that (ii) PII uridylylation depends exclusively on UT/UR in response to glutamine levels (Schumacher *et al*, 2013; Jiang *et al*, 2007; van Heeswijk *et al*, 2013). We validated both assumptions by conducting similar time course experiments as above and showing that in a *glnK* knock-out strain GS adenylylation was not significantly changed and that in a UT/UR (*glnD*) knock-out PII was 100% non-uridylylated (Fig. Supp. 3).

## DISCUSSION

An organism’s survival depends on its ability to sense and interact appropriately with its environment. At the cellular level such processing of information is marshalled by molecular interaction networks. These take up cues from the environment and the cell’s physiological state, and transduce this information to the relevant cellular response machinery, which includes regulators of protein activity and transcriptional activators, as well as their down-stream targets. The precise way in which biological information is being processed is still largely unknown for most important signalling networks: while the connections for many signal transduction and stress-response pathways are known, or at least partly known (Huvet *et al*, 2011), the details and the dynamics of information flow are only beginning to be understood (Mc Mahon *et al*, 2014) (Cheong *et al*, 2011; Uda *et al*, 2013). In particular, the ability of signal transduction networks to distinguish between quantitatively different signals, such as for instance ambient levels of different nutrients, is typically not well understood. Even for the better understood signal transduction networks (Clausznitzer *et al*, 2010; Kollmann *et al*, 2005) we have only sketchy knowledge of the way information is processed, or how information is mapped by the molecular machineries in order to allow the organism to deal with changing environmental conditions.

The channel capacities that we found for the nitrogen sensing system are high, especially when compared to other systems that have recently been analyzed in a similar framework (Tkacik *et al*, 2008; 2009; Cheong *et al*, 2011; Kollmann *et al*, 2005). (These studies were performed on single cells, with a number of advantages, but also possible disadvantages, e.g. fluorescent reporters may introduce additional sources of biochemical variability and substantial measurement noise) As techniques for single cell measurements of PTMs or metabolite abundances are not (yet) available, we took the approach of making population measurements, and then reconstructing noise levels based on biophysical predictions of stochasticity in biochemical signalling.) However, they are consistent with recent theoretical predictions e.g. (Ziv *et al*, 2007; Sarpeshkar, 2014), and provide an important indication that currently available lower bounds on information capacity of biochemical signalling pathways may be substantially underestimated. Most previous studies have investigated signalling capacities that act at the level of protein expression or nuclear translocation, while the PII-GS PTM signalling system considered here directly acts on the metabolic level by regulating GS enzyme activity. The high cellular abundances of PII and GS are partly responsible for the high information processing capacity that we found; and there are good biological reasons for why this system should perform with such a high fidelity (Endres & Wingreen, 2008; Kentner & Sourjik, 2009): *E. coli* processes the environmental nitrogen state with high accuracy so that metabolic fluxes can be finely tuned to cellular needs and nutrient availability. For microbes (as opposed to previous studies on isolated mammalian cells), there is a very direct link between sensing levels of environmental nutrients and evolutionary fitness (Klumpp & Hwa, 2014), especially given the large numbers of genomes and rapid generation times of bacteria, and so even subtle improvements in the fidelity of signal transduction will afford a fitness advantage (Lynch, 2012).

Nitrogen assimilation through GS is a high-throughput metabolic pathway, with approximately one million glutamate + NH_4_^+^ conversions into glutamine per cell per second during exponential growth (Schumacher *et al*, 2013). Here we have focussed on the two key components of the nitrogen sensing machinery. We find that the response tracks the availability of ambient nitrogen faithfully, continuously and with high fidelity – the machinery is not simply switched ON or OFF in response to changes in nitrogen availability, but is instead poised in a state of optimal sensitivity: at nitrogen concentrations that are encountered frequently, the concentrations of the nitrogen currencies GLN and αKG are reflected accurately by the activity levels of GS and PII. This is, of course, entirely reasonable as a less accurate regulation of such a major metabolic pathway would carry a fitness penalty.

The accuracy of nitrogen sensing appears to be reduced as the abundances of GLN and αKG enter regimes that should be rarely if ever encountered. This also makes perfect sense from both an information-theoretical as well as an evolutionary perspective. GS, PII and their associated (de-) adenylylating and (de-)uridylylating enzymes form the information channels along which input (GLN and αKG states) is transmitted, processed and returned as output (GS-A and PII-U). Shannon’s noisy channel coding theorem (Cover & Thomas, 2012) tells us that the channel capacity is the maximum information that can flow through a channel. This will be precisely the case if the most frequent signals are transmitted with high fidelity (Fig. 4), and the Fisher information strikes the necessary balance between input frequency and information processing fidelity. This outcome is intimately linked to the evolutionary relevance: life-history theory tells us that competition between individuals will be fiercest under commonly encountered conditions. Adaptation to rarely encountered environments is rarely beneficial (Stumpf *et al*, 2002) as the required trade-offs typically entail poor adaptation under other, more frequently encountered conditions; in this sense the Fisher information can indeed be the subject of selection (Frank, 2009). Signalling systems such as the GS-PII system can also form the basis for rationally designed (Barnes *et al*, 2011) biological sensing and computing devices, which naturally implement efficient inference procedures.

Our theoretical methodology establishes a general and computationally efficient framework to analyze information processing in biochemical circuits. It enables us to quantify the information capacity and, more importantly, to understand how reliable molecular signalling processes can be. Information processing is thus of fundamental importance for bacterial survival. Being able to rationally manipulate or adapt such sensing machines will also have implications for biotechnology and agriculture as nitrogen metabolism is crucial to e.g. plant growth.

## References

1. Aebersold R, Burlingame AL & Bradshaw RA (2013) Western blots versus selected reaction monitoring assays: time to turn the tables?. Mol. Cell Proteomics 12: 2381–2382

2. Barnes CP, Silk D, Sheng X & Stumpf MPH (2011) Bayesian design of synthetic biological systems. Proc. Natl. Acad. Sci. U.S.A. 108: 15190–15195

3. Bruggeman FJ, Boogerd FC & Westerhoff HV (2005) The multifarious short-term regulation of ammonium assimilation of Escherichia coli: dissection using an in silico replica. 272: 1965–1985

4. Brunel N & Nadal JP (1998) Mutual information, Fisher information, and population coding. 10: 1731–1757

5. Bueno R, Pahel G & Magasanik B (1985) Role of glnB and glnD gene products in regulation of the glnALG operon of Escherichia coli. J. Bacteriol. 164: 816–822

6. Cheong R, Rhee A, Wang CJ, Nemenman I & Levchenko A (2011) Information Transduction Capacity of Noisy Biochemical Signaling Networks. Science 334: 354–358

7. Clausznitzer D, Oleksiuk O, Løvdok L, Sourjik V & Endres RG (2010) Chemotactic response and adaptation dynamics in Escherichia coli. 6: e1000784

8. Kustu S, Hirschman J, Burton D, Jelesko J & Meeks JC (1984) Covalent modification of bacterial glutamine synthetase: physiological significance. 197: 309–317

9. Cover TM & Thomas JA (2012) Elements of Information Theory John Wiley & Sons

10. Daniel R, Rubens JR, Sarpeshkar R & Lu TK (2013) Synthetic analog computation in living cells. 497: 619–623

11. Endres RG & Wingreen NS (2008) Accuracy of direct gradient sensing by single cells. PNAS 105: 15749–15754

12. Endres RG & Wingreen NS (2009) Maximum likelihood and the single receptor. 103: 158101

13. Frank SA (2009) Natural selection maximizes Fisher information. 22: 231–244

14. Hart Y, Madar D, Yuan J, Bren A, Mayo AE, Rabinowitz JD & Alon U (2011) Robust control of nitrogen assimilation by a bifunctional enzyme in E. coli. 41: 117–127

15. Huvet M, Stumpf MPH, Toni T, Sheng X, Thorne TW, Jovanovic G, Engl C, Buck M, Pinney JW & Stumpf MP (2011) The evolution of the phage shock protein response system: interplay between protein function, genomic organization, and system function. Mol Biol Evol 28: 1141–1155

16. Jiang P, Mayo AE & Ninfa AJ (2007) Escherichia coli glutamine synthetase adenylyltransferase (ATase, EC 2.7.7.49): kinetic characterization of regulation by PII, PII-UMP, glutamine, and alpha-ketoglutarate. 46: 4133–4146

17. Jiang P, Peliska JA & Ninfa AJ (1998) Enzymological characterization of the signal-transducing uridylyltransferase/uridylyl-removing enzyme (EC 2.7.7.59) of Escherichia coli and its interaction with the PII protein. 37: 12782–12794

18. Kentner D & Sourjik V (2009) Dynamic map of protein interactions in the Escherichia coli chemotaxis pathway. Molecular Systems Biology 5:

19. Khoury GA, Baliban RC & Floudas CA (2011) Proteome-wide post-translational modification statistics: frequency analysis and curation of the swiss-prot database. Sci Rep 1:

20. Kim M, Zhang Z, Okano H, Yan D, Groisman A & Hwa T (2012) Need-based activation of ammonium uptake in Escherichia coli. Molecular Systems Biology 8: 616

21. Klumpp S & Hwa T (2014) Bacterial growth: global effects on gene expression, growth feedback and proteome partition. Curr. Opin. Biotechnol. 28C: 96–102

22. Kollmann M, Løvdok L, Bartholomé K, Timmer J & Sourjik V (2005) Design principles of a bacterial signalling network. 438: 504–507

23. Komorowski M, Costa MJ, Rand DA & Stumpf MPH (2011) Sensitivity, robustness, and identifiability in stochastic chemical kinetics models. Proc. Natl. Acad. Sci. U.S.A. 108: 8645–8650

24. Komorowski M, Finkenstädt B, Harper CV & Rand DA (2009) Bayesian inference of biochemical kinetic parameters using the linear noise approximation. BMC Bioinformatics 10:

25. Komorowski M, Zurauskiene J & Stumpf MPH (2012) StochSens--Matlab package for sensitivity analysis of stochastic chemical systems. Bioinformatics 28: 731–733

26. Libby E, Perkins TJ & Swain PS (2007) Noisy information processing through transcriptional regulation. Proc. Natl. Acad. Sci. U.S.A. 104: 7151–7156

27. Liepe J, Barnes CP, Kirk P, Filippi S, Toni T & Stumpf MPH (2014) A framework for parameter estimation and model selection from experimental data in systems biology using approximate Bayesian computation. Nat Protoc 9: 439–456

28. Lodeiro A & Melgarejo A (2008) Robustness in Escherichia coli glutamate and glutamine synthesis studied by a kinetic model. J Biol Phys 34: 91–106

29. Lynch M (2012) Evolutionary layering and the limits to cellular perfection. PNAS 109: 18851–18856

30. Ma H, Boogerd FC & Goryanin I (2009) Modelling nitrogen assimilation of Escherichia coli at low ammonium concentration. J. Biotechnol. 144: 175–183

31. Mc Mahon SS, Sim A, Filippi S, Johnson R, Liepe J, Smith D, Stumpf MP & Stumpf MPH (2014) Information theory and signal transduction systems: from molecular information processing to network inference. Seminars in Cell & Developmental Biology 35: 98–108

32. Mehta P, Goyal S, Long T, Bassler BL & Wingreen NS (2009) Information processing and signal integration in bacterial quorum sensing. Molecular Systems Biology 5:

33. Ninfa AJ, Jiang P, Atkinson MR & Peliska JA (2000) Integration of antagonistic signals in the regulation of nitrogen assimilation in Escherichia coli. Curr. Top. Cell. Regul. 36: 31–75

34. Okano H, Hwa T, Lenz P & Yan D (2010) Reversible adenylylation of glutamine synthetase is dynamically counterbalanced during steady-state growth of Escherichia coli. J. Mol. Biol. 404: 522–536

35. Picotti P & Aebersold R (2012) Selected reaction monitoring-based proteomics: workflows, potential, pitfalls and future directions. 9: 555–566

36. Sarpeshkar R (1998) Analog versus digital: extrapolating from electronics to neurobiology. 10: 1601–1638

37. Sarpeshkar R (2014) Analog synthetic biology. Philos Trans A Math Phys Eng Sci. 372: 20130110–20130110

38. Schumacher J, Behrends V, Pan Z, Brown DR, Heydenreich F, Lewis MR, Bennett MH, Razzaghi B, Komorowski M, Barahona M, Stumpf MPH, Wigneshweraraj S, Bundy JG & Buck M (2013) Nitrogen and carbon status are integrated at the transcriptional level by the nitrogen regulator NtrC in vivo. MBio 4: e00881–13

39. Selimkhanov J, Taylor B, Yao J, Pilko A, Albeck J, Hoffmann A, Tsimring L & Wollman R (2014) Accurate information transmission through dynamic biochemical signaling networks. Science 346: 1370–1373

40. Stumpf MPH, Laidlaw Z & Jansen VAA (2002) Herpes viruses hedge their bets 99: 15234–15237

41. Tkacik G, Callan CG & Bialek W (2008) Information flow and optimization in transcriptional regulation. PNAS 105: 12265–12270

42. Tkacik G, Walczak AM & Bialek W (2009) Optimizing information flow in small genetic networks. Phys. Rev. E (3). 80: 031920

43. Toni T, Welch D, Strelkowa N, Ipsen A & Stumpf MPH (2009) Approximate Bayesian computation scheme for parameter inference and model selection in dynamical systems. J. R. Soc. Interface 6: 187–202

44. Uda S, Saito TH, Kudo T, Kokaji T, Tsuchiya T, Kubota H, Komori Y, Ozaki Y-I & Kuroda S (2013) Robustness and Compensation of Information Transmission of Signaling Pathways. Science 341: 558–561

45. van Heeswijk WC, Westerhoff HV & Boogerd FC (2013) Nitrogen assimilation in Escherichia coli: putting molecular data into a systems perspective. Microbiol. Mol. Biol. Rev. 77: 628–695

46. Van Kampen NG (2011) Stochastic Processes in Physics and Chemistry Elsevier

47. Wallace EWJ, Gillespie DT, Sanft KR & Petzold LR (2012) Linear noise approximation is valid over limited times for any chemical system that is sufficiently large. IET Syst Biol 6: 102–115

48. Wedel A, Weiss DS, Popham D, Dröge P & Kustu S (1990) A bacterial enhancer functions to tether a transcriptional activator near a promoter. Science 248: 486–490

49. Yuan J, Doucette CD, Fowler WU, Feng X-J, Piazza M, Rabitz HA, Wingreen NS & Rabinowitz JD (2009) Metabolomics-driven quantitative analysis of ammonia assimilation in E. coli. 5: 302

50. Zimmer DP, Soupene E, Lee HL, Wendisch VF, Khodursky AB, Peter BJ, Bender RA & Kustu S (2000) Nitrogen regulatory protein C-controlled genes of Escherichia coli: scavenging as a defense against nitrogen limitation. 97: 14674–14679

51. Ziv E, Nemenman I & Wiggins CH (2007) Optimal signal processing in small stochastic biochemical networks. PLoS ONE 2: e1077

